# Human research islet cell culture outcomes at the Alberta Diabetes Institute IsletCore

**DOI:** 10.1101/2024.06.18.599388

**Authors:** James G Lyon, Alice LJ Carr, Nancy P Smith, Braulio Marfil-Garza, Aliya F Spigelman, Austin Bautista, Doug O’Gorman, Tatsuya Kin, AM James Shapiro, Peter A Senior, Patrick E MacDonald

## Abstract

Human islets from deceased organ donors have made important contributions to our understanding of pancreatic endocrine function and continue to be an important resource for research studies aimed at understanding, treating, and preventing diabetes. Understanding the impacts of isolation and culture upon the yield of human islets for research is important for planning research studies and islet distribution to distant laboratories. Here we examine islet isolation and cell culture outcomes at the Alberta Diabetes Institute (ADI) IsletCore (n=197). Research-focused isolations typically have a lower yield of islet equivalents (IEQ), with a median of 252,876 IEQ, but a higher purity (median 85%) than clinically-focused isolations before culture. The median recovery of IEQs after culture was 75%, suggesting some loss. This was associated with a shift towards smaller islet particles, indicating possible islet fragmentation, and occurred within 24 hours with no further loss after longer periods of culture (up to 136 hours). No overall change in stimulation index as a measure of islet function was seen with culture time. These findings were replicated in a representative cohort of clinical islet preparations from the Clinical Islet Transplant Program at the University of Alberta. Thus, loss of islets occurs within 24 hours of isolation and there is no further impact of extended culture prior to islet distribution for research.

## Introduction

Since the development of the first large scale purification protocols,^1^ human pancreatic islets have become a valuable resource of tissue for research studies and in clinical transplant programs. They are used in a diverse range of research projects related to islet morphology, cell proliferation, genomics, insulin and glucagon secretion, fuel-induced toxicity, transcription factor regulation, transplantation, and many other aspects of endocrine physiology and diabetes.^2,3^ Furthermore, deceased donor islets are used in clinical transplant programs for the management of type 1 diabetes.^4,5^ Several clinical islet transplant programs exist,^6^ that may provide tissue for research either following ‘unsuccessful’ clinical islet isolations of insufficient yield for transplant or by performing research-specific isolations.^7,8^ Distribution of such research islets are managed either in-house, as with the Alberta Diabetes Institute (ADI) IsletCore (www.isletcore.ca), or *via* one of several networks, such as the Integrated Islet Distribution Program. Relatively few academic islet isolation programs focus solely on research-specific islet isolations.^7^

At the ADI IsletCore (www.isletcore.ca) research-specific islet isolations are performed, with written informed research consent, using pancreata that have not been accepted for clinical use. Key features of this research-only program include the non-GMP nature of isolations performed at ADI IsletCore and a focus on islet purity over total yield. This is largely accomplished by combining fewer density gradient fractions to increase islet isolation purity. Following isolation, research islets are typically cultured between 1-5 days prior to shipment to the end user. This is usually done to avoid shipment over weekends, where delivery to recipient laboratories is often not possible. It is important to understand if this extended culture time contributes to islet loss in culture prior to shipment. In the University of Alberta Clinical Islet Transplant Program (CITP), a long-standing transplantation program working in parallel to the ADI IsletCore, factors such as organ preservation, cold ischemic time, islet size, and preparation purity have been shown to impact islet recovery after ∼20 hours.^9^ Others have shown impacts of tissue seeding density,^10^ preparation purity,^11^ and temperature.^12^ Here we examine isolation outcomes at the ADI IsletCore and factors that influence islet loss in culture overnight and with extended pre-shipment culture. We demonstrate the role for islet size and possible fragmentation to loss of islets in culture but show that most changes occur within 24 hours of culture, with little further impact of extended pre-shipment culture time. Key findings are validated in a representative, separate, cohort of islet preparations from the University of Alberta CITP.

## Materials and Methods

### Human research pancreata and islet isolation

We included 197 non-diabetic deceased human pancreatic donors from which the islets were isolated by the ADI IsletCore between September 2016 and January 2023. Of these, most donors were the neurological determination of death donors (82%) with the remainder following donation after circulatory death (28%). These research-specific islets were isolated and cultured using standard culture techniques,^13^ followed by the quantification of the islets before and after culture.

Briefly, all pancreata were perfused in a controlled manner with an enzyme solution of collagenase (Liberase MTF; Roche Diagnostics; or CIzyme Collagnease HA; VitaCyte, LLC or Collagenase Gold 800; VitaCyte, LLC), with most islet isolations using Vitacyte Collagenase Gold (Vitacyte LLC; 69%), and a non-specific protease (Thermolysin; Roche Diagnostics or BP Protease; VitaCyte, LLC) via the pancreatic duct. Islets were separated by mechanical dissociation using a Ricordi Islet Isolator (Biorep) and purified using a continuous gradient of a polymer based separating solution (Biochrom or Lympholite) in an apheresis system (model 2991 COBE; Terumo BCT, Inc.). Gradient fractions containing the higher purity of islet tissue were combined with the goal of optimum islet purity.

Human islet isolation and the use of human islets in research was approved by the Human Research Ethics Board at the University of Alberta (Pro00001754, Pro00013094). All donors’ families gave informed consent for use of pancreatic tissue for research.

### Human islet culture

The isolated islets were placed in culture prior to distribution or experimentation in CMRL 1066 (Corning) supplemented with 0.5% BSA (Equitech-Bio), 1% insulin-transferrin-selenium (Corning), 100 U/mL penicillin/streptomycin (Life Technologies), and L-glutamine (Sigma-Aldrich). All isolated islets were cultured in non-treated petri dishes between 4 to 136 hours (median 33 hours (IQR 18;62.00)) at 22^°^C with 5% CO_2_at a typical seeding density of 225 IEQ/cm^2^.

### Islet quantity, purity, and functionality assessment

Prior to culture, isolations were assessed in duplicate, and purity and percentage of trapped islets were determined.^14^ By using the dithizone contrast dye, the Islets were quantified prior to and following culture by islet equivalents (IEQ), the standard unit for reporting variations in the volume of islets correct to 150 µm diameter,^15^ and islet particle number (IPN). Islet diameter was determined, and islets were categorized into the following bins: 50-100, 101-150, 151-200,201-250, 251-300,301-350, >351 µm. Islet particle index (IPI), which is an indication for the size of islets relative to a standard 150 µm diameter IEQs, was calculated as IPN divided by number of IEQs. Islets were assayed for total cellular insulin (Meso Scale Discovery, Rockville, MD, USA) and DNA (Quant-iT™ PicoGreen ® dsDNA, Molecular Probes, Eugene, OR, USA). Prior to and following culture the islets were assessed for percent purity and portion trapped in contaminating exocrine tissue. By using the dithizone contrast dye,^15^ the islets were visually assessed by generated images obtained with a stereoscope with a ring light to obtain a black contrast background (Olympus MVX10, with a DP72 camera). Insulin and DNA content for quality control,^16^ and glucose-stimulated insulin secretion,^17^ were assessed as described.

### Glucose stimulated insulin secretion

Glucose-stimulated insulin secretion measurements were performed at 37^°^ C in Krebs-Ringer buffer (KRB) (in millimoles: NaCl 115; KCl 5; NaHCO_3_ 24; CaCl_2_ 2.5; MgCl_2_ 1; HEPES 10; 0.1% BSA, pH7.4) with glucose concentrations as noted.^18^ Triplicate groups of 15 hand-picked islets were preincubated for 2 hours with 2.8 mM glucose KRB. Islets were subsequently incubated for 1 hour in 2.8 mM glucose KRB, followed by 1 hour of stimulation in 16.7 mM glucose KRB. The supernatants were collected and the insulin content was extracted from the islet pellet using acid-ethanol. Samples were stored at −20°C and assayed for insulin via electrochemiluminescence (Meso Scale Discovery SA). Outliers in the values of insulin content at 2.8 mM and 16.7 mM were identified as values more than 3 times the median absolute deviation away from the median of each respective insulin content. Stimulation index was calculated as *(insulin secretion at 16*.*7 mM / insulin secretion at 2*.*8 mM* for each replicate of the triplicate that did not contain an outlier in insulin content, and then averaged across the triplicate.

### Replication cohort

A set of 172 clinical islet isolations from the University of Alberta CITP were sampled from isolations performed over the same time period as above, that experienced culture. Isolation and quantification protocols of the CITP can be found elsewhere.^19^

### Statistical methods

Analyses was performed using R statistical software version 4.3.0 (Foundation for Statistical Computing, Vienna, Austria). Data are presented as median and interquartile range (IQR). Summary statistics were generated using the gtsummary package (version 1.7.1).^20^ Comparisons of descriptive characteristics were made by Wilcoxon rank sum test for unpaired tests or Wilcoxon signed rank test with continuity correction for paired tests (pre-culture to post-culture). Comparison across more than two unpaired groups was assessed by Kruskal-Wallis test. Subsequent post-hoc analysis for pairwise comparisons was conducted using Wilcoxon rank sum test adjusted using Bonferroni correction. We repeated the assessment of factors influencing ‘major islet loss’ (defined as >20% loss) in the ADI IsletCore, that has been previously demonstrated in the CITP ^9^, using univariate and multivariable logistic regression. Covariates included the continuous variables: donor age, body mass index (BMI), cold ischemia time, pancreas weight, digestion time, preculture purity, preculture trapped islets, preculture IPI and preculture IPN and the dichotomous variables: Sex and culture time of >24 hrs. Significance was tested at a level of P<0.05.

## Results

### Donor and isolation characteristics

Donors included in this study were adult (>18 years of age) research pancreas donors (n=197). Properties of the donors and isolations are shown in **Table 1**. Compared to reports from clinical islet isolation programs,^21–24^ including the University of Alberta CITP,^9,25,26^ islet yield for these research-specific isolations was lower while purity was higher. This is expected given the focus of these isolations on combining fewer density gradient fractions of higher purity.

**Table 1.** Donor, organ and isolation characteristics and pre- and post-culture quantification at the Alberta Diabetes Institute IsletCore.

| Characteristic |  | ADI IsletCore,<br>N = 197* |
| --- | --- | --- |
| Donor<br>Characteristics | Age (years) | 51 (39; 60) |
|  | Sex (Male) | 123 (62%) |
|  | BMI (kg/m<sup>2</sup>) | 26.9 (24.2; 30.3) |
|  | HbA1c (%) | 5.50 (5.10; 5.70) |
|  | Unknown | 25 |
|  | Blood Glucose (mmol/l) | 9.8 (8.5; 11.5) |
|  | Unknown | 15 |
| Organ/Isolation<br>Characteristics | Cold ischemia time (hours) | 13.0 (10.5; 16.0) |
|  | Pancreas weight (g) | 90 (76; 106) |
|  | Unknown | 0 |
|  | Collected tissue (g) | 38 (27; 50) |
|  | Unknown | 0 |
|  | Digestion time (mins) | 17.0 (15.0; 19.0) |
|  | Undigested tissue (g) | 13 (9; 19) |
|  | Unknown | 0 |
|  | Culture time (hours) | 33 (18; 62) |
|  | Purity (Pre-culture) (%) | 85 (75; 90) |
|  | Trapped islets (Pre-culture) (%) | 0 (0; 0) |
|  | Unknown | 0 |
| Pre-culture<br>quantification | Total islet equivalents (IEQ) | 252,876<br>(174,147; 367,152) |
|  | Islet particle number (IPN) | 237,000<br>(162,000; 299,000) |
|  | Islet particle index (IPI) | 1.11 (0.83; 1.52) |
| Post-culture<br>quantification | Total islet equivalents (IEQ) | 195,272<br>(128,535; 255,659) |
|  | Islet particle number (IPN) | 217,000<br>(149,000; 285,000) |
|  | Islet particle index (IPI) | 0.85 (0.66; 1.13) |
|  | Recovery of IEQs (%) | 75 (64; 91) |

### IEQ loss during culture

IEQ loss in culture results in fewer available to ship for research. Median recovery of IEQs post-culture was 75%, with a significant decrease post-culture in the median total number of IEQs (p<0.0001), IPN (p=0.008) and DNA content (p<0.0001) (**Figure 1A,B,C**; **Suppl Table 1**). Median insulin content did not significantly change, however, post-culture (**Figure 1D, Suppl Table 1**).

**Figure 1.**
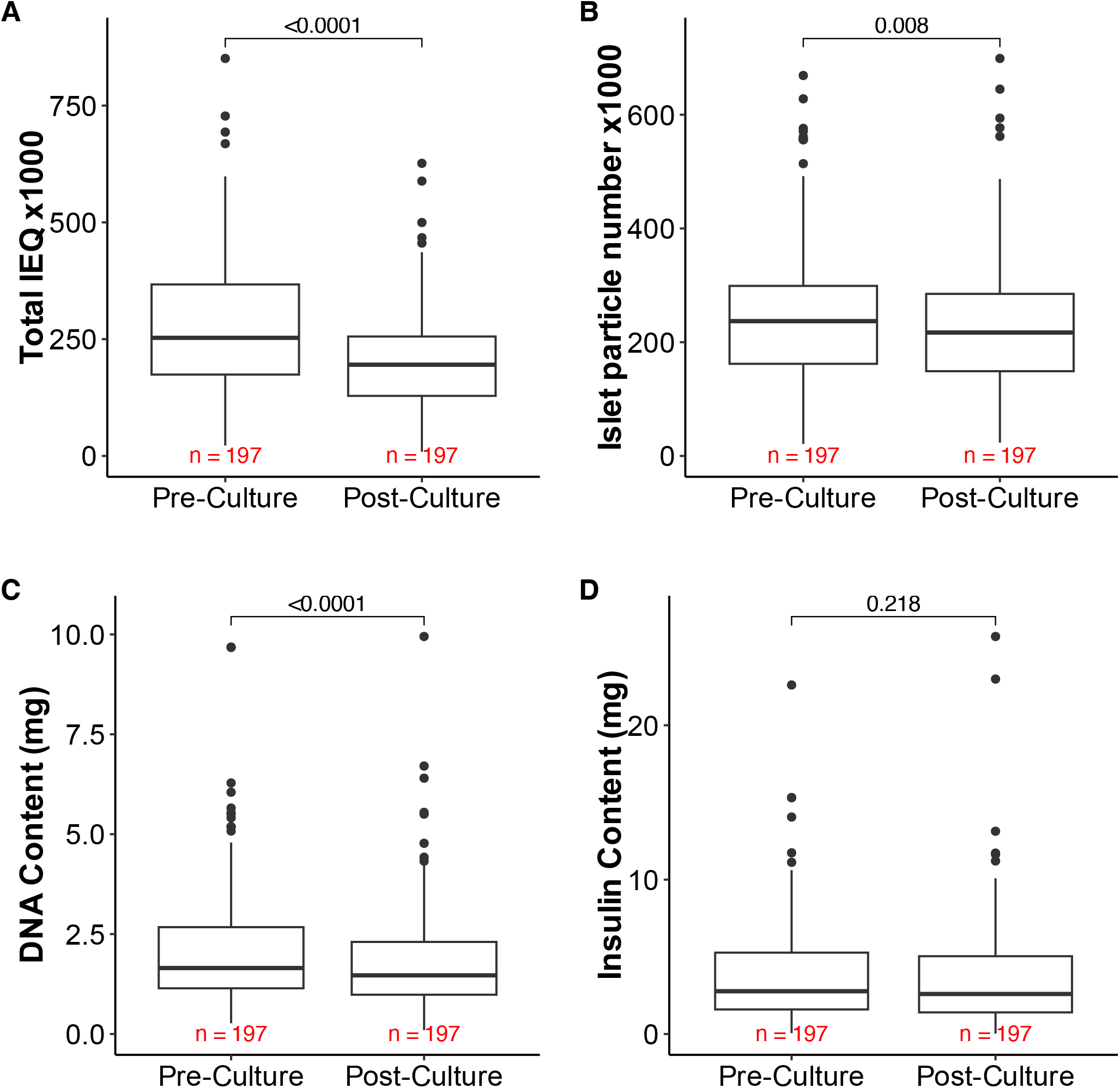
Comparison of pre- and post-culture: Total number of Islet Equivalents (IEQ) (A), Islet particle number (IPN) (B), DNA content (C) and Insulin content (D) in **Alberta Diabetes Institute IsletCore**.

Islet size (indicated by IPI values) decreased post-culture (p<0.0001) (**Figure 2A, Suppl Table 1**), and there was a significant increase in the proportions of smaller categorized islets of diameter 50-100 µm (p<0.0001) whereas the proportions of larger diameter categorized islets decreased post-culture (p<0.0001) for all categories of diameter larger than 151 µm (**Figure 2B**). This suggests the loss of larger islets during culture, and an increase in smaller islets that could represent ‘islet fragmentation’ leading to an overall loss of IEQ counts (particularly since fragments smaller than 50 µm diameter will not be counted).

**Figure 2.**
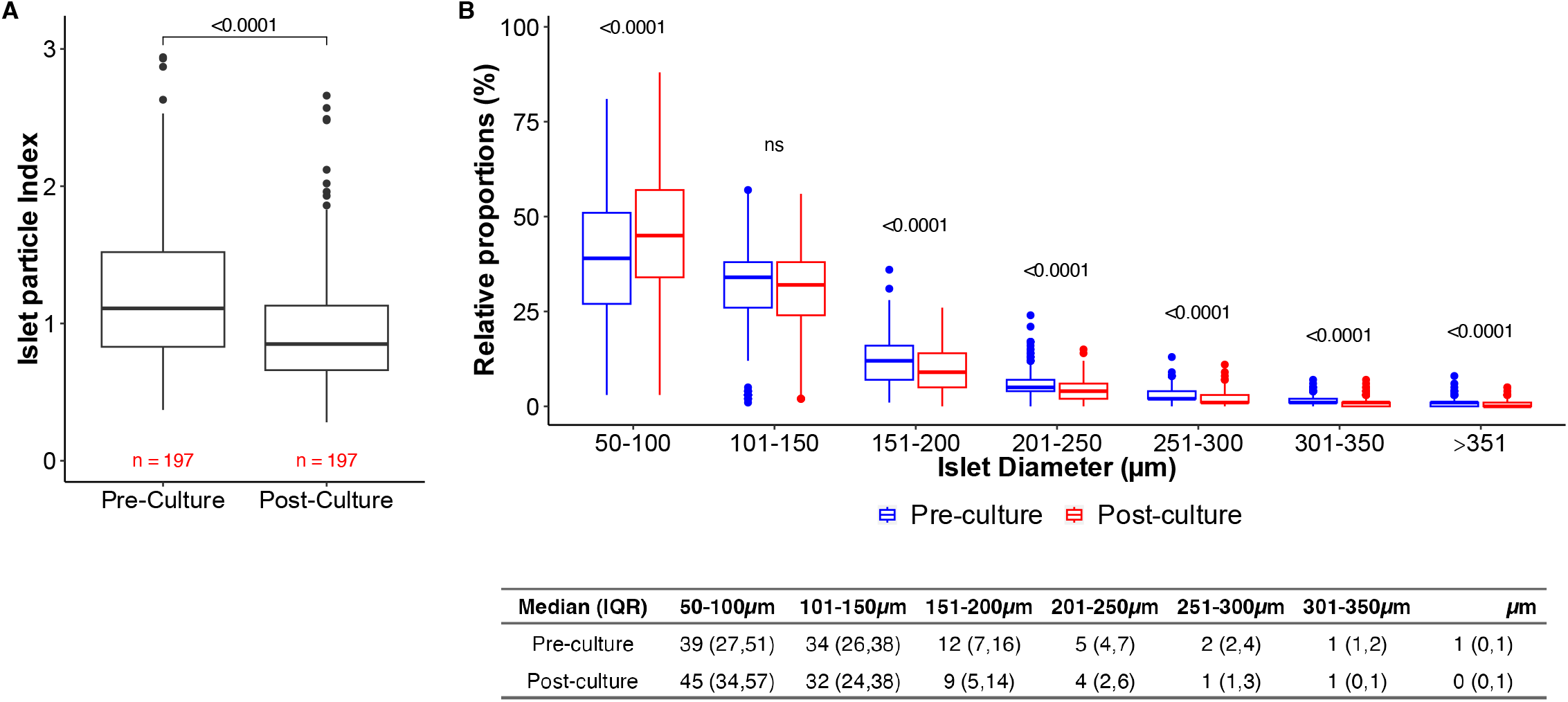
Comparison from pre- to post-culture of islet particle index (A) and the relative proportions of islet diameter categories, with the median and interquartile range of the proportions displayed in the adjoined table (B) in **Alberta Diabetes Institute IsletCore**.

### A proportion of isolations increased in islet particle number post-culture

Although ‘islet fragmentation’ should increase the overall number of islet particles in culture, we find that IPN decreased post-culture (**Suppl Table 1**). This is likely because any resulting dithizone-positive fragments <50 µm diameter would not be counted. However, at an individual level a large minority of isolations (42.6%; n=84/197) showed an increase in IPN post-culture. These isolations had significantly larger IPI to start at pre-culture (p=0.0008), compared to isolations whose IPN decreased or had no change (**Suppl Table 2**). Culture time, total IEQs pre-culture, percentage of trapped islets pre-culture and insulin and DNA content were not significantly different between isolations with an increased IPN post-culture and isolations whose IPN decreased or had no change (**Suppl Table 2**). Of note, isolations with a percentage increase in IPN post-culture had higher percentage recovery of IEQs post-culture (**Suppl Table 2**), likely because in these cases more of the fragments remained in the countable range (i.e. >50 µm diameter). Conversely, isolations where IPN is not seen to change (or decreases) may have a more dramatic loss of IEQ as fragments fall below the countable range.

### Larger islet particle Index, higher pancreas weight and older age were factors influencing major islet loss

An effect of islet size on ‘major islet loss’ (defined as >20% loss) in culture was demonstrated previously within the Clinical Islet Transplant Program.^9^ By this criteria we identified that 114/197 (58%) IsletCore isolations experienced major islet loss. In univariate multivariable logistic regression, while controlling for other covariates and their interaction, we confirmed that IPI had substantial effect on major islet loss, with isolations with larger sized islets being 3.26 [95% CI 1.59, 7.05; p=0.002] more likely to experience major IEQ loss compared to those with smaller islets (**Table 2**). Additionally, we found older age and heavier pancreas weight to be modest predictors of significance for islet loss (Age: 1.03 [95% CI 1.00,1.05] p=0.024 and pancreas weight: 1.02 [1.00,1.03] p=0.033) (**Table 2**). However no significant effect was found for BMI, cold ischemia time, digestion time, preculture purity, preculture trapped islets, preculture IPN, sex and culture time of >24 hrs (**Table 2**).

**Table 2.** Odds ratios, 95% confidence intervals and significance level for univariate and multivariable logistic regression, in addition to each covariates summary metrics, in the assessment of factors influencing ‘major islet loss’ (defined as >20% loss) in the ADI IsletCore.

| Characteristic | N=197* | Univariate |  |  | Multivariable |  |  |
| --- | --- | --- | --- | --- | --- | --- | --- |
|  |  | OR | 95% CI | p-value | OR | 95% CI | p-value |
| Age (years) | 51 (39, 60) | 1.02 | 1.00, 1.04 | **0.031** | 1.03 | 1.00, 1.05 | **0.024** |
| Male Sex (Reference: Female) | 123 (62%) | 1.28 | 0.72, 2.30 | 0.4 | 0.81 | 0.40, 1.61 | 0.6 |
| BMI (kg/m<sup>2</sup>) | 26.9 (24.2, 30.3) | 1.03 | 0.97, 1.09 | 0.4 | 0.97 | 0.91, 1.04 | 0.4 |
| Cold ischemia time (hours) | 13.0 (10.5, 16.0) | 1.01 | 0.96, 1.07 | 0.7 | 1.01 | 0.95, 1.08 | 0.7 |
| Pancreas Weight (g) | 90 (76, 106) | 1.01 | 1.00, 1.02 | **0.026** | 1.02 | 1.00, 1.03 | **0.033** |
| Digest time (min) | 17.0 (15.0, 19.0) | 0.97 | 0.89, 1.05 | 0.5 | 0.97 | 0.88, 1.07 | 0.5 |
| Purity (preculture) (%) | 85 (75, 90) | 1.00 | 0.98, 1.01 | >0.9 | 0.99 | 0.97, 1.01 | 0.4 |
| Trapped islets (preculture) (%) | 0.0 (0.0, 0.0) | 0.99 | 0.97, 1.02 | 0.5 | 0.99 | 0.96, 1.02 | 0.4 |
| Islet particle index (preculture) | 1.11 (0.83, 1.52) | 2.04 | 1.14, 3.79 | **0.019** | 3.26 | 1.59, 7.05 | **0.002** |
| Islet particle number, x1,000 (preculture) | 237,000 (162,000, 299,000) | 1.00 | 1.00, 1.00 | 0.6 | 1.00 | 1.00, 1.00 | 0.2 |
| Culture time >24 hours (h) (Reference: ≤ 24 hrs) | 115 (58%) | 1.34 | 0.76, 2.39 | 0.3 | 1.47 | 0.78, 2.78 | 0.2 |

\*Median (IQR)

### Culture time beyond 24 hours has no impact on the recovery or function of islets

No overall trend between post-culture recovery of IEQs and culture time was observed p=0.72 (**Figure 3A**), with a marginally significant trend observed with stimulation index (p=0.049) (**Figure 3B**). However, upon post-hoc testing, there was no significant difference in the post-culture recovery of IEQs or stimulation index in isolations cultured for ≤24 hours, as compared to isolations cultured for longer, up to a maximum of 136 hours. The median recovery of IEQs in isolations cultured for ≤24 hours was 77.5% (IQR 65.25,91.75) and median stimulation index was 6.09 (IQR 3.62,9.68).

**Figure 3.**
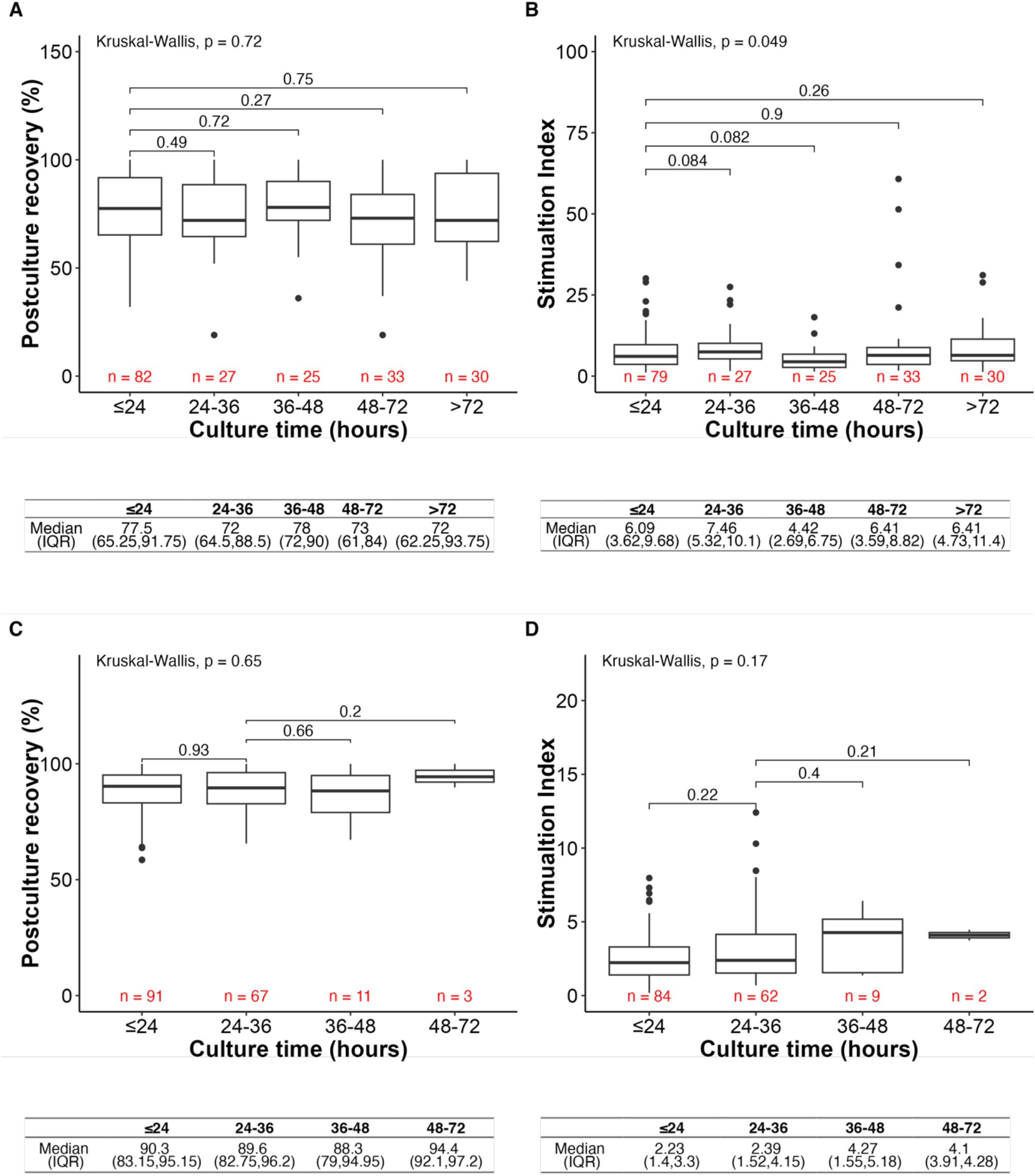
Relationship between culture time with recovery of IEQs post-culture (A) and Stimulation Index from Glucose Stimulated Insulin Secretion (B) **Alberta Diabetes Institute IsletCore**, comparative to the **University of Alberta Hospital Clinical Islet Transplant program** (C and D respectively). The post-hoc within pairs testing p-value shown is from Wilcox rank sum adjusted with Bonferroni correction. Median and interquartile range for recovery and stimulation index at each culture time category are displayed below each respective figure.

### Cell culture outcomes in a replication cohort

We assessed key cell culture outcomes in a sample of 172 islet isolations performed at the University of Alberta CITP over the same time frame. Although recovery of IEQs after culture was significantly higher at 90% (83, 96) vs. 75 (64, 91), p<0.0001), we confirmed decreasing IEQs, IPI and IPN post-culture (p<0.0001; p<0.0001; p<0.0001) (**Suppl Table 3**). As with the research-specific isolations from the ADI IsletCore, 43.3% (n=52/172) of isolations at the CITP increased IPN post-culture. Unlike the ADI IsletCore, in the CITP sample we did not see that these isolations had significant differences in purity pre-culture (p=0.31) or in islets size pre-culture (p=0.15), compared to isolations whose IPN decreased or had no change (**Suppl Table 4**). Similarly to the ADI IsletCore however, in the CITP sample we observed the total IEQs preculture were significantly lower in isolations where a percentage increase in IPN post-culture (p=0.039) occurred. In addition, these isolations had a higher percentage recovery of IEQs post-culture (**Suppl Table 4**).

Similarly to the ADI IsletCore, in the CITP there was no overall trend across culture time in the post-culture recovery of IEQs (p=0.65) or simulation index (p=0.17) (stimulation index present for 157/172 isolations). Post hoc testing confirmed there was no difference in these rates of recovery or stimulation index for isolations cultured for ≤24 hours, as compared to isolations cultured for up to a maximum of 72 hours (**Figure 3C,D**).

## Discussion

Human islets for research have become an important component of the diabetes research ecosystem,^3^ and efforts to improve quality control and reporting are aimed at improving research reproducibility in the field.^27–31^ Because research islets are often cultured prior to shipment to investigators, it is important to determine the impact of such culture on isolation yield and function. Since pre-shipment culture time may be required for logistic reasons, it is important to understand whether waiting 1 day or 4 days can impact the material available for research studies. We have found that, despite an initial islet loss in culture, there seems to be no further detrimental loss or obvious impairment in function or insulin content beyond 24 hours. This gives us some reassurance when needing to wait a few days before islet shipment to researchers.

It is thought that larger islets are often lost in culture due to hypoxia-induced cell death^32^ and possible fragmentation. Indeed, human islet fragmentation is increased by induced hypoxia in culture.^33^ We observe a shift from larger to smaller islet sizes in culture, with an overall reduction in IPN which would equate to increase in what was previously called the ‘islet fragmentation index’ (which is the converse of islet particle index, calculated as IPN/IEQ or 1/IPI).^34^ We find that 47% of isolations increase in IPN after culture, which may be explained by the fragmentation of islets. Why this is not observed in all preparations is likely due to either the complete loss of some islet fragments, or fragmentation to the point where they are not counted in the IEQ calculation bins. This could also explain why preparations with a larger IPI tend to increase in IPN after culture, while preparations with a smaller IPI do not. In short, when preparations with a smaller initial islet size undergo some degree of fragmentation this results in fragments that are below the threshold for counting, whereas the fragmentation of larger islets are more likely to result in additional countable particles (although some must clearly be lost, as IPI correlates with loss of IEQs).

Finally, we importantly demonstrate that islet loss in our program and the replication cohort occurs within the first 24 hours of culture and is then stable up to 5 days. In addition, we show no change in insulin content before and after culture. This is important given our need to often culture islets before shipment to researchers (or, in the case of the CITP, to culture prior to transplant). This, along with the stability of the insulin secretory phenotype, gives us some reassurance. One important caveat though is that these studies represent a post hoc analysis of outcomes, rather than a longitudinal study of islet morphology and function. But it is important to note that this does not rule the possibility that islets change in culture. An early stress response following isolation has been demonstrated,^35–37^ which may decrease with a day or two in culture. Indeed, we recently demonstrated that human islet transcriptome is dramatically altered in culture but that the proteome is much less affected (in preparation), which may explain why we observed no impact on insulin secretion in the present analysis. Although we previously reported that culture of human islets from a limited number of donors for greater than 4 days reduces stimulation index,^38^ comparison with the current results is difficult since our longest culture group was greater than or equal to 3 days. It seems likely that prolonged culture would eventually have a detrimental impact on islet function, although our primary concern here was the impact within our normal pre-shipment culture time.

Thus, we report significant human islet loss after in culture after isolation in line with previous reports. This results from islet fragmentation, with a shift from larger to smaller islet particles, and a subsequent loss of counted islet fragments when they fall below the 50µm diameter cutoff. Importantly, the majority of this occurs within 24 hours and that additional culture within the normal pre-shipment window for the ADI IsletCore seems to have no further detrimental effect.

## Data sharing

Data from ADI IsletCore on donor and isolation outcomes analyzed here are freely available at www.humanislets.com.

## Acknowledgements

Human islets for research were provided by the Alberta Diabetes Institute IsletCore at the University of Alberta in Edmonton (http://www.bcell.org/adi-isletcore.html) with the assistance of the Human Organ Procurement and Exchange program, Trillium Gift of Life Network, and other Canadian organ donation organizations. Islet isolation was approved by the Human Research Ethics Board at the University of Alberta (Pro00013094). All donors’ families gave informed consent for the use of pancreatic tissue in research.

## Funding

Work was funded by grants from the Canadian Institutes of Health, JDRF and Diabetes Canada. AMJS holds a Canada Research Chair (Tier 1) in Transplantation Surgery and Regenerative Medicine. PAS is the Charles A Allard Chair in Diabetes Research and is supported by the Academic Medicine and Health Services Program. PEM holds a Canada Research Chair (Tier 1) in Islet Biology.

## Declaration of interests

AMJS serves as a consultant to Vertex Inc, Aspect Biosystems Inc. and Betalin Ltd, and is a co-inventor for a patent on TNFRSF25-mediated treatments of immune diseases and disorders (PCT/US2020/053085) and for a Cellular Transplant Site-Device-less technology (US 14/863541, CA.286512)”

PAS has received speaker fees from Abbott, Dexcom, GSK, Insulet, LMC, Novo Nordisk, Vertex; and consulting fees from Abbott, Bayer, Dexcom, GSK, Insulet, Novo Nordisk, Sanofi, Vertex, Ypsomed.

All other authors have no competing interests to declare.

## Figure Legends

**Figure 1. Post-culture loss of islets at the Alberta Diabetes Institute IsletCore**

Comparison of pre- and post-culture total islet equivalents (IEQ) (**A**), islet particle number (IPN) (**B**), insulin content (**C**), and DNA content (**D**) in preparations from the Alberta Diabetes Institute IsletCore. Data shows median and interquartile ranges.

**Figure 2. Loss of larger islets and gain of smaller islets post-culture**

Comparison from pre- to post-culture of islet particle index (IPI) (**A**) and the relative proportions of islet diameter categories (**B**) from the Alberta Diabetes Institute IsletCore. Data shows median and interquartile ranges.

**Figure 3. Islet loss occurs within 24-hours, and then is unchanged for greater than 72-hours**

Relationship between culture time with recovery of islet equivalents (IEQs) post-culture and stimulation index (SI) of islet preparations from the Alberta Diabetes Institute IsletCore (**A,B**) and in a replication cohort from the University of Alberta Clinical Islet Transplant Program (**C,D**). The post-hoc within pairs testing p-value shown is from Wilcox rank sum adjusted with Bonferroni correction. Median and interquartile range are shown.

## Notes

https://www.humanislets.com

