## Supplementary Materials for "Human research islet cell culture outcomes at the Alberta Diabetes Institute IsletCore"

**Supplementary Table 1** Comparison of pre- and post-culture islet quantification and size in Alberta Diabetes Institute IsletCore.

| **Characteristic** | **Pre-culture**, N = 197* | **Post-culture**, N = 197* | **p-value**† |
|---|---|---|---|
| Total IEQs | 252,876 (174,147, 367,152) | 195,272 (128,535, 255,659) | **<0.0001** |
| Islet particle number | 237,000 (162,000, 299,000) | 217,000 (149,000, 285,000) | **0.008** |
| Insulin Content (mg) | 2.77 (1.59, 5.27) | 2.59 (1.40, 5.04) | 0.2 |
| DNA Content (mg) | 1.65 (1.14, 2.67) | 1.47 (0.98, 2.30) | **<0.0001** |
| Islet particle index | 1.11 (0.83, 1.52) | 0.85 (0.66, 1.13) | **<0.0001** |

*Median (IQR)
†Wilcoxon signed rank test with continuity correction

**Supplementary Table 2** Comparison of pre-culture characteristics and IEQ recovery rates between isolations that had a post-culture percentage increase in islet particle number (IPN) and those that had no change or a decrease, in Alberta Diabetes Institute IsletCore.

| **Characteristic** | **Increased IPN**, N = 84* | **No change or decreased IPN**, N = 113* | **p-value**† |
|---|---|---|---|
| Culture time (hours) | 31 (18, 45) | 36 (18, 65) | 0.1 |
| Post-culture Recovery of IEQs (%) | 85 (72, 95) | 72 (58, 82) | **<0.0001** |
| Purity (preculture) (%) | 90 (75, 90) | 80 (75, 90) | **0.018** |
| Islet Particle Index (preculture) | 1.23 (1.02, 1.59) | 1.01 (0.79, 1.49) | **0.0008** |
| Trapped islets (preculture) (%) | 0.0 (0.0, 0.0) | 0.0 (0.0, 5.0) | 0.1 |
| Total IEQs (preculture) | 262,986 (199,640, 377,111) | 251,357 (165,573, 338,714) | 0.2 |
| DNA Content (preculture) (mg) | 1.64 (1.15, 2.69) | 1.69 (1.14, 2.67) | 0.7 |
| Insulin Content (preculture) (mg) | 3.11 (1.56, 5.32) | 2.59 (1.65, 5.02) | 0.7 |

*Median (IQR)

†Wilcoxon rank sum test

**Supplementary Table 3** Comparison of the available pre- and post-culture quantification and size characteristics in University of Alberta Hospital Clinical Transplant Program.

| **Characteristic** | **Pre-culture**, N = 172* | **Post-culture**, N = 172* | **p-value**† |
|---|---|---|---|
| Total IEQs | 544,083 (444,099, 635,385) | 465,117 (392,604, 569,426) | **<0.0001** |
| Islet particle number | 440,922 (350,750, 535,673) | 407,374 (328,371, 515,937) | **<0.0001** |
| Islet particle index | 1.24 (1.09, 1.44) | 1.19 (1.01, 1.37) | **<0.0001** |

*Median (IQR)
†Wilcoxon signed rank test with continuity correction

**Supplementary Table 4** Comparison of the available pre-culture characteristics and IEQ recovery rates between isolations that had a percentage increase in islet particle number post-culture and those that had no change or a decrease, in The University of Alberta Hospital Clinical Transplant Program.

| **Characteristic** | **Increased IPN**, N = 52* | **No change or decreased IPN**, N = 120* | **p-value**† |
|---|---|---|---|
| Culture time (hours) | 19 (16, 31) | 25 (19, 32) | 0.07 |
| Post-culture Recovery of IEQs (%) | 95 (89, 100) | 87 (80, 94) | **<0.0001** |
| Purity (preculture) (%) | 50 (45, 55) | 50 (45, 60) | 0.31 |
| Islet Particle Index (preculture) | 1.28 (1.12, 1.48) | 1.21 (1.05, 1.43) | 0.15 |
| Trapped islets (preculture) (%) | 6 (3, 13) | 5 (2, 8) | 0.1335 |
| Total IEQs (preculture) | 501,882 (410,426, 614,551) | 560,076 (461,937, 647,112) | **0.039** |

*Median (IQR)

†Wilcoxon signed rank test with continuity correction
